# Knockdown-resistance (*kdr*) mutations in Indian *Aedes aegypti* populations: lack of recombination among haplotypes bearing V1016G, F1534C, and F1534L *kdr* alleles

**DOI:** 10.1101/2024.12.11.627862

**Authors:** Taranjeet Kaur, Rajababu S. Kushwah, Sabyasachi Pradhan, Manoj K. Das, Madhavinadha P Kona, Anushrita, David Weetman, Rajnikant Dixit, Om P. Singh

## Abstract

**Background:** Knockdown resistance (*kdr*) mutations in the voltage-gated sodium channel (VGSC) gene are a key mechanism of insecticide resistance in mosquitoes. In Asian *Aedes aegypti* populations two main VGSC haplogroups with *kdr* mutations have been identified: one carrying the F1534C mutation and another with V1016G and/or S989P mutations. Functional studies have demonstrated that these three mutations on a single haplotype confer up to a 1100-fold increase in pyrethroid resistance, underscoring the need to monitor these triple mutations within distinct populations. This study investigates the prevalence of *kdr* mutations in Indian populations and explores the linkage association between these mutations and two distinct conserved types of introns located between exons 20 and 21.

**Methods:** *Ae. aegypti* specimens collected from eight different locations were genotyped for *kdr* alleles and intron (between exons 20 and 21) haplotypes using PCR-based assays. Representative samples underwent DNA sequencing of VGSC regions.

**Results:** Five *kdr* mutations namely S989P, V1016G, T1520I, F1534C, and F1534L, were identified, each exhibiting varying distribution and frequencies across different geographical regions. Two distinct and stably-diverged intron haplotypes, designated as intron-A and intron-B, were identified between exons 20 and 21. Seven haplotypes, including two wild-type variants, were observed among Indian populations. The *kdr*-bearing haplotypes can be classified into three distinct haplogroups: haplogroup G (V1016G with/or without S989P and with intron-A), haplogroup L (F1534L and intron-A), and haplogroup C (F1534C with/or without T1520I and with intron-B). Importantly, no evidence of recombination within Indian populations was detected among these three haplogroups.

**Conclusions:** Five *kdr* mutations were identified in the VGSC of Indian *Ae. aegypti* populations, each showing a definitive linkage with one of the two types of intron haplotypes. The lack of recombination among haplogroups bearing 1016G with 989P, 1534C and 1534L mutations suggests that the most potent insecticide resistance haplotype, bearing the triple *kdr* mutation, is currently absent. This finding has significant operational implications, as it may indicate that current vector control measures remain effective against these populations, potentially delaying the emergence of highly resistant phenotypes

## Background

*Aedes aegypti* serves as a vector for several human arboviral infections including dengue virus (DENV), chikungunya virus (CHIKV), yellow fever virus (YFV), and Zika virus (ZIKV). Controlling these arboviral infections relies mainly on vector control measures. Pyrethroids, a commonly used group of insecticides for disease vector control, are favoured due to their low toxicity to mammals and rapid insect knockdown action. These insecticides act on insects by binding their open sodium channels and modifying the channel-gating kinetics by inhibiting the inactivation resulting in prolonged opening of the sodium channel and paralysis, and eventual insect death. Despite their efficacy, the effectiveness of pyrethroid-based control of *Ae. aegypti* is increasingly challenged by the escalating development of pyrethroid resistance worldwide [1], including in India [2-4]. Therefore, achieving effective management of resistance necessitates the ongoing monitoring of insecticide resistance patterns, a comprehensive understanding of the underlying mechanisms, and the identification of key markers for surveillance.

Knockdown resistance (*kdr*) is one of the mechanisms of insecticide resistance against DDT and pyrethroids which is due to the mutation/s in the voltage-gated sodium channel (VGSC) - the target site of action for DDT and pyrethroids. These mutations render the VGSC less susceptible to DDT and pyrethroids. The most commonly reported *kdr* mutation, L1014F/S, is not present in *Ae. aegypti* due to codon constraints [5], where a minimum of two SNPs are required in the codon to replace wild type Leucine with either resistant Phenylalanine or Serine amino acids. In *Ae. aegypti*, at least 23 mutations have been reported in the VGSC [6,7, 8]. Among these, amino acid variants at the 1534 and 1016 VGSC residues are the most well-documented mutations, and their role in pyrethroid resistance has been functionally validated [9-10]. The distribution of *kdr* alleles show significant variation across different regions of the world. There are two alternative *kdr* mutations reported at two residues V1016 (G/I) and F1534 (C/L) which exhibited distinct geographical distributions. Mutation V1016I is reported from the Americas, multiple African countries and Iran [11, 12, 13], while the alternative mutation V1016G (and linked mutation S989P) is reported from countries in Southeast Asia and Saudi Arabia [14] but has not been reported from countries in the Americas [11], except in Panama [15]. The T1520I is reported only from three Asian countries, India [7-16], Myanmar [17-18] and Laos [13].

Co-occurrence of multiple *kdr* mutations often with strong linkage associations have been reported, for example, S989P [3, 14, 19,20,21] and/or D1763Y [22-23] have been found in conjunction with V1016G while T1520I is associated with F1534C [3, 18, 24]. These linkages may have an additive or multiplicative effect on insecticide resistance levels [25, 26] or may have a compensatory effect to overcome the deleterious effect of mutation on the fitness of the mosquito [3, 27, 28]. Linkage of *kdr* mutations has also been shown with two distinct intron-haplotypes linking exon 20 and 21 of the VGSC, named intron A and B [29, 30, 31]. Functional analysis of various sodium channel types by Hirata et al. [26], as expressed in *Xenopus* oocytes, revealed that S989P+V1016G and F1534C individually reduced their sensitivity to permethrin by 100- and 25-fold, respectively. However, a VGSC with the S989P+V1016G+F1534C triple mutation present on same haplotype, reduced sensitivity by 1100-fold against permethrin and by 90-fold against deltamethrin. Thus, a single recombination could lead to critical failure of a pyrethroid-based vector control programme; therefore monitoring the occurrence of the triple *kdr* mutation in populations is viewed as crucial [26]. The present study was undertaken to monitor *kdr* mutations in Indian populations and screen for the presence of triple mutant *kdr* mutations (S989P+V1016G+F1534C), if any. In addition, the study also sought to establish linkage associations of different *kdr* mutations with the two intron types in Indian populations.

## Material and methods

### Mosquito sampling

*Aedes aegypti* immatures were collected from natural breeding habitats across various regions of India (Figure 1), namely New Delhi (28.58° N, 77.05° E), Bhopal, Madhya Pradesh (23.26° N, 77.41° E), Khandwa, Madhya Pradesh (21.83° N, 76.35° E), Ranchi, Jharkhand (23.35° N, 85.30° E), Raipur, Chhattisgarh (21.25° N, 81.63° E), Kolkata, West Bengal (22.57° N, 88.36° E), and Chennai, Tamil Nadu (13.08° N, 80.27° E). Late-stage larvae (III-IV instars) and pupae were sampled from breeding sites and transferred to plastic bowls containing water until pupation. Larvae were fed with a mixture of ground dog biscuit and yeast in a ratio of 3:1. Pupae were subsequently moved to bowls with water and placed in mosquito cages. Inside the mosquito cage, a cotton pad soaked in a 10% aqueous glucose solution served as a food source for the mosquitoes. Upon emergence, the mosquitoes were anesthetized using diethyl ether and identified morphologically under a stereo dissecting microscope. Morphologically confirmed *Ae. aegypti* specimens were stored individually in microcentrifuge tubes, containing dehydrated silica gel wrapped in a small piece of paper. The collected mosquitoes were transported to the laboratory in Delhi and stored at -20 °C until DNA isolation.

**Figure 1.**
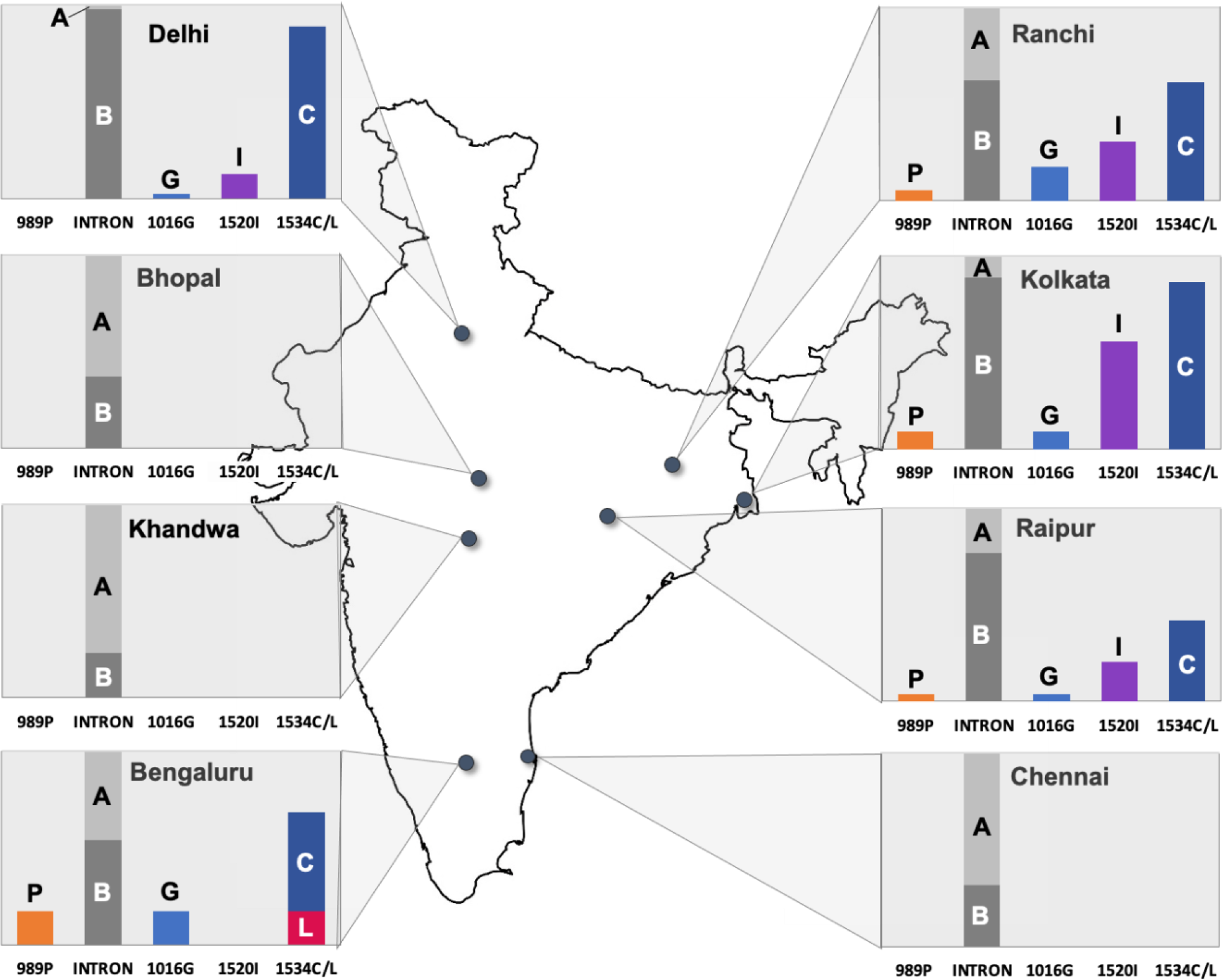
Frequencies of *kdr* mutant alleles (P=989P, G=1016G, I=1520I, C=1534C, L=1534L) and intron types (A= intron type A, B=intron type B) in different populations of India.

### DNA isolation

The DNA was isolated from individual mosquitoes using the method of Black & Duteau [32]. Prior to DNA isolation from female mosquitoes, the last three segments of the abdomen were excised to eliminate the spermatheca, which may contain sperms from male mating partners. The elution of DNA was performed in 200 μL of TE buffer and stored at 4 °C for immediate use or at -20 °C for prolonged storage. Additionally, DNA samples isolated from field-collected mosquitoes in Bengaluru in a previous study by Kushwah et al. [7] were also incorporated into this present study.

### Genotyping of *kdr* alleles

Genotyping of *kdr* alleles was conducted on DNA extracted from individual mosquitoes using PCR-based techniques developed by Kushwah et al. [3, 7]. The list of primers is provided in Table 1. Briefly, for genotyping of domain III-*kdr* mutations i.e., F1534C, F1534L and T1520I, a single PCR product was amplified using primers AekdrF and AekdrR followed by restriction digestion of the PCR product with three different restriction enzymes, i.e., *BsaB*I (for T1520I), *Ssi*I (for F1534C), and *Eco88*I (for F1534L)—each in separate 0.5 mL microcentrifuge tubes. For genotyping of S989P and V1016G *kdr* mutations, two independent allele-specific PCRs were performed using common flanking primers AedIIF and AedIIR along with two allele-specific primers, i.e., PPF and SSR for S989 alleles and VVF and GGR for V1016 alleles (Table 1) following the methodology described by Kushwah et al. [7]. All PCRs were carried out using Hot Start DNA polymerase (DreamTaq, ThermoFisher).

**Table 1.**
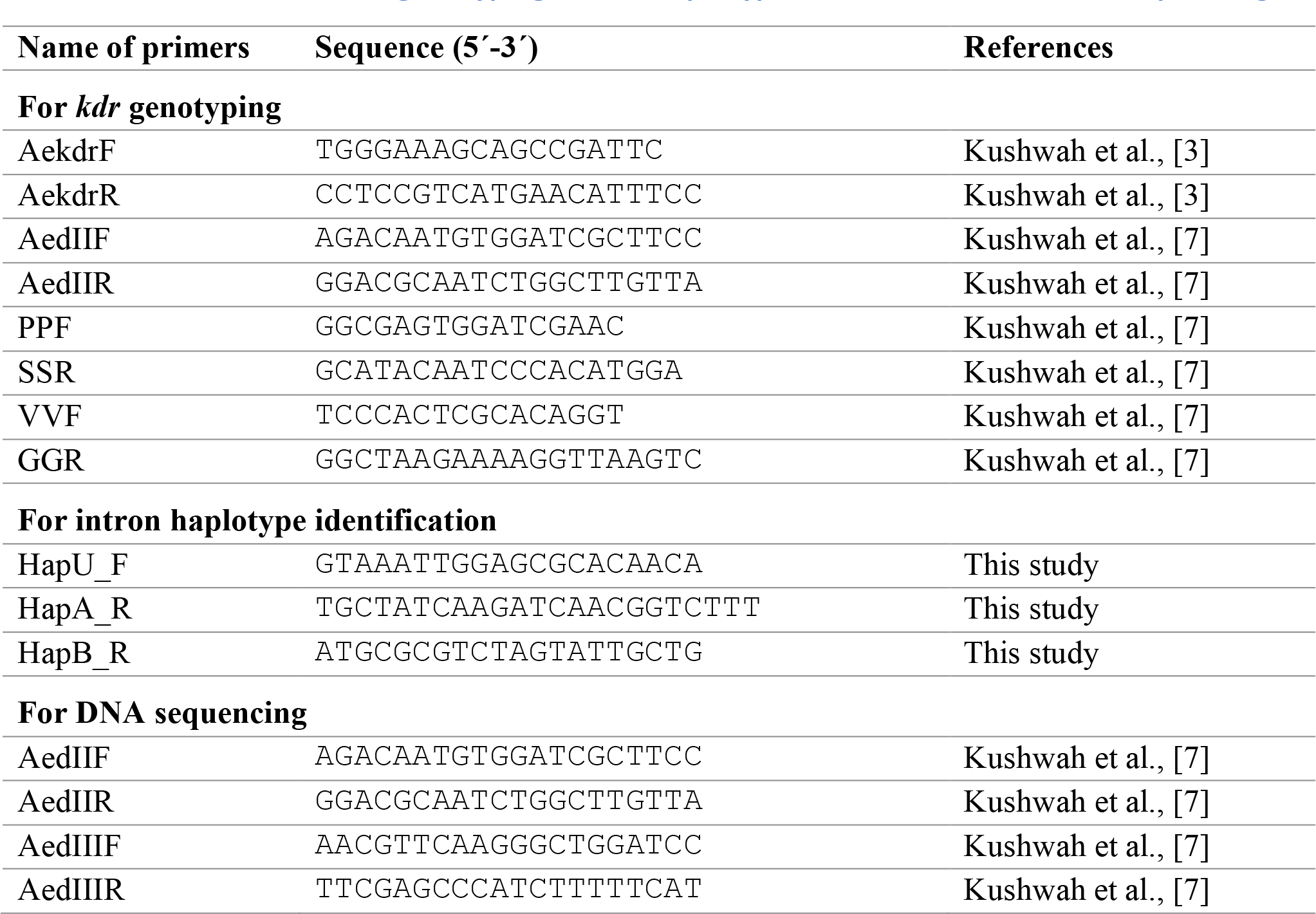
Primers used for *kdr* genotyping, intron haplotype identification, and DNA sequencing.

### Identification of intron-haplotype using PCR

As observed by earlier workers [23, 30, 33, 34], two distinct intron haplotypes (connecting exons 20 and 21) were observed in domain II of the VGSC with a high degree of nucleotide and size polymorphism, which were named A and B following Martins et al. [30]. For the identification of intron types, a multiplex PCR strategy was developed. A universal forward primer HapU_F (Table 1) was designed from a conserved region of the intron. Two intron haplotype-specific reverse primers, HapA_R and HapB_R (Table 1), were designed which were specific for haplotypes A and B, respectively. The intron haplotype-specific primers were designed in such a way that at least five nucleotides at the 3’ end of each primer had mismatches with the non-target haplotype to ensure that there was no non-specific extension. Multiplex PCR amplification was performed using 2X DreamTaq Hot Start DNA polymerase (ThermoFisher) in a 15 µl reaction volume. Each reaction contained 0.2 µM of each primer, i.e., HapU_F, HapA_R and HapB_R. The thermal cycling program included an initial denaturation at 95 °C for 3 minutes, followed by 35 cycles consisting of denaturation at 95 °C for 15 seconds, annealing at 58 °C for 15 seconds, and extension at 72 °C for 30 seconds. A final extension was carried out at 72 °C for 7 minutes. PCR products were separated on a 2% agarose gel and visualized under UV light using a gel documentation system. The diagnostic amplicon sizes for haplotypes A and B were, 81 and 128 bp, respectively (Figure 2).

**Figure 2.**
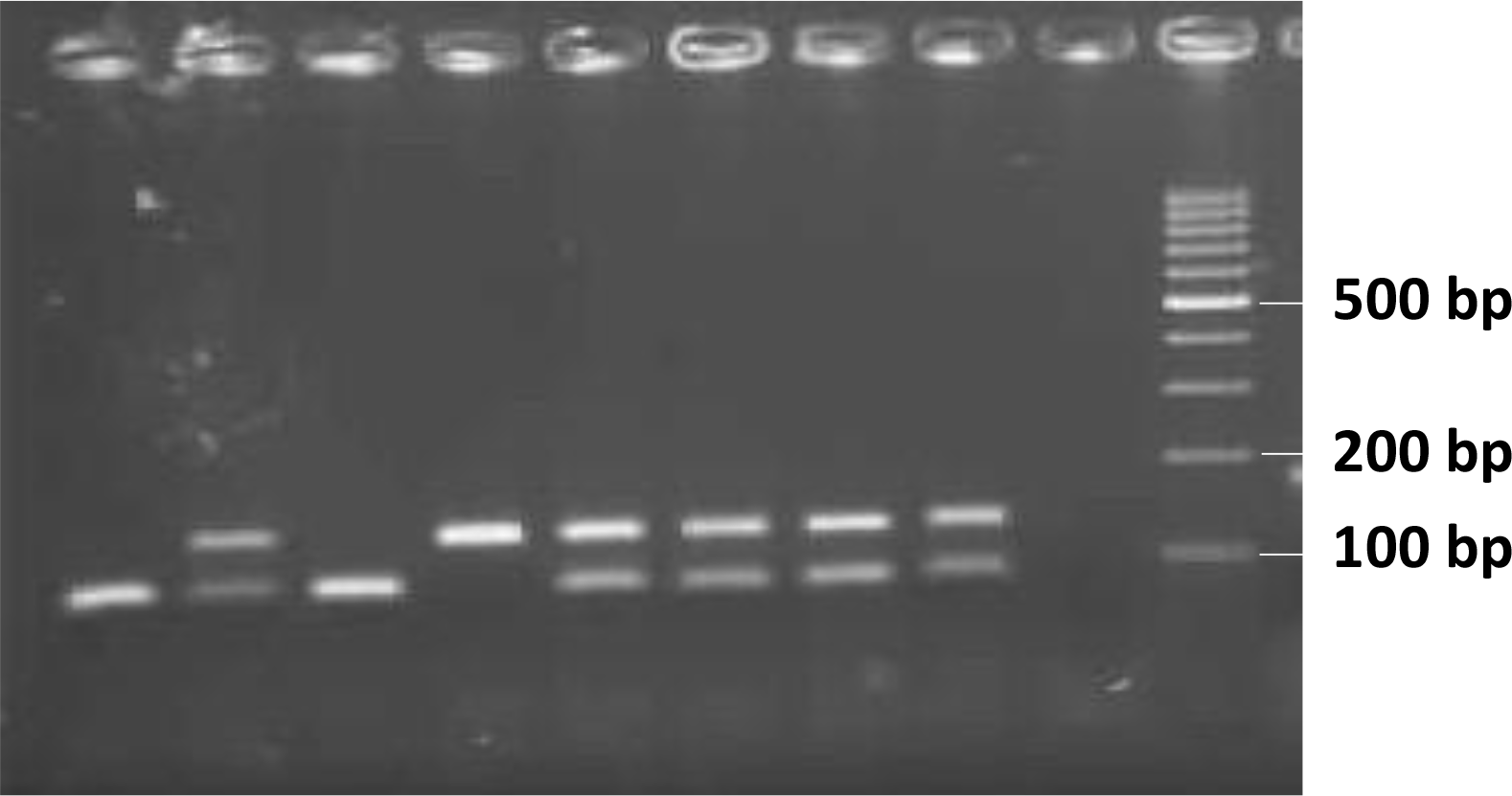
Gel photograph of PCR used for genotyping intron haplotypes. An 81 bp amplicon represents intron type A, while a 128 bp amplicon represents intron type B. Lanes 1 to 8 are test samples, lane 9 is negative control (without DNA) and lane 10 is 100 bp DNA ladder.

### Haplotype phasing

Estimation of haplotype definitions and frequencies based on genotyping results of loci S989, V1016, T1520, F1534 and intron types, was performed on pooled data using the Gibbs sampling strategy implemented in Arlequin ver 3.5 [35]. Since this method relies on homozygote data for reliable haplotype phasing, and some populations lacked homozygotes at certain loci, the data were pooled across populations for the analysis.

### DNA sequencing

Sequencing was performed on DNA samples isolated from individual mosquitoes for partial sections of domains II and III. The PCR amplifications for domain II were carried out using primers AedIIF and AedIIR, while for domain III, primers AedIIIF and AedIIIR from Table 1 were employed, following the method by Kushwah et al. [7]. PCR products underwent ExoSAP-IT (Thermo Fisher) treatment for the removal of primers and dNTPs and were sequenced using ABI BigDye Terminator v3.2 (ThermoFisher). The sequences were analyzed using Finch TV ver. 1.5.0. The number of samples sequenced for both domains II and III were 79 (mostly homozygous for intron types) and 68 respectively.

## Results

Five *kdr* mutations were identified, S989P and V1016G in domain II, and T1520, F1534C, and F1534L in domain III of the VGSC, each exhibiting varying geographical distribution and frequencies (Figure 1; Table 2). Mutations were recorded in a total of five populations, i.e., Delhi, Ranchi, Raipur, Kolkata and Bengaluru while no *kdr* mutations were found in three populations, i.e., Bhopal, Khandwa and Chennai. The most dominant mutation present in all five populations having *kdr* mutations was F1534C with allelic frequencies ranging from 42–89%. The second most common mutation was T1520I present in four populations, i.e., Delhi, Raipur, Ranchi, and Kolkata with allele frequencies ranging from 13–56%. The mutation V1016G was present in these five populations with low frequencies ranging from 2-18%. The mutation S989P was present in Ranchi, Raipur, Kolkata and Bengaluru with frequencies ranging from 2–18%. The mutation F1534L was only detected in Bengaluru, as reported previously [7].

**Table 2.**
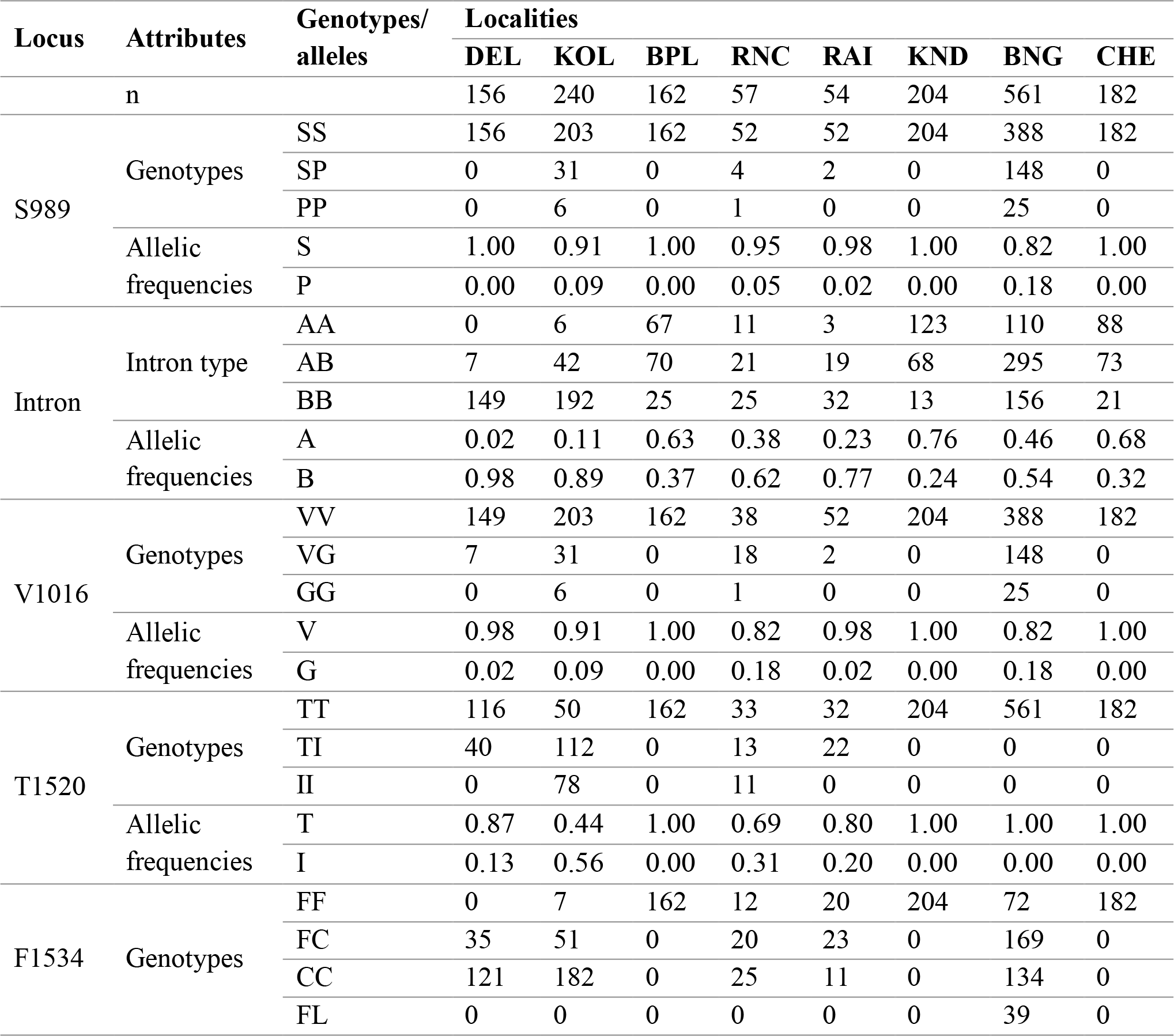
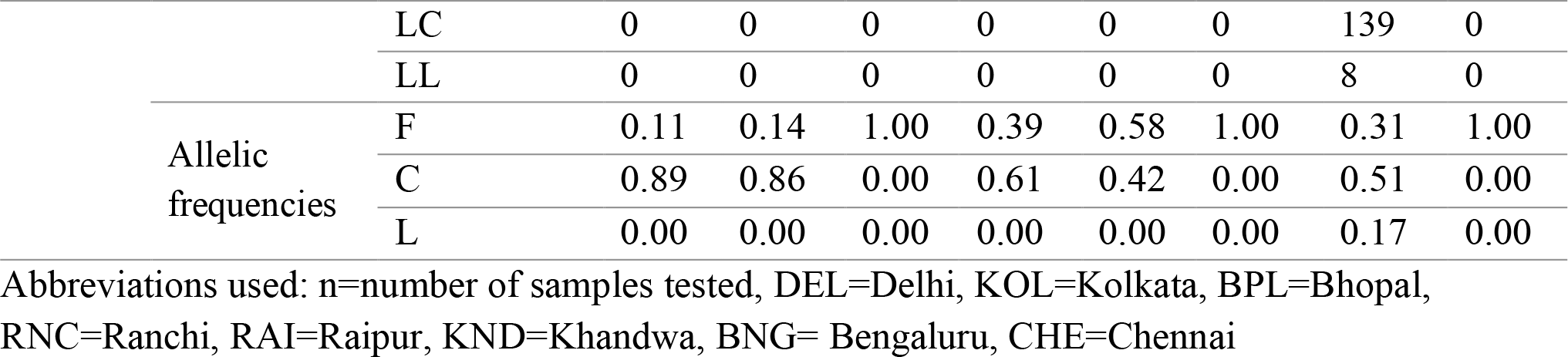
*kdr*-genotypes and intron types scored in different populations.

RNC=Ranchi, RAI=Raipur, KND=Khandwa, BNG= Bengaluru, CHE=Chennai DNA sequencing of domain IIS6 among the samples which were homozygous for specific intron haplotype alleles, revealed the presence of only two intron types, belonging to either intron-haplotype A and B. The two types of introns were highly conserved, as evidenced by comparison of the DNA sequences of 70 samples which were homozygotes for intron type (21 intron type A and 49 intron type B) from Ranchi, Kolkata, and Bengaluru with those from Fan et al. (2020) [36]. Intron A matched the DNA sequences of haplotypes V4 (S989/V1016), V8 (989P/1016G), and V9 (S989/1016I), while intron B was identical to haplotype V2 [36]. Genotyping of samples using intron-specific primers revealed that the proportion of these two intron haplotypes varied in different populations. The frequencies of intron-A ranged between 2 and 76% and of intron-B ranged between 24 and 98%. The proportion of intron B was highest in populations having *kdr* mutations (54–98%) while it was lowest in populations with no *kdr* (23–37%).

Phasing revealed the presence of a total of seven haplotypes including two wild haplotypes considering the polymorphisms at locus S989, intron type (connecting exon 20 and 21), V1016, T1520, and F1534. The number of samples with different genotype combinations including intron type in each population is shown in Table 3. The definition of haplotypes and their numbers is shown in Table 4. The *kdr* alleles showed clear linkage association with other *kdr* alleles and intron types. The 1016G mutant was inked to intron-A and was present with or without 989P. The 1534C and 1534L mutants were linked to intron-B and former was present with or without 1520I. The five haplotypes with *kdr* allele(s) can be categorized into three haplogroups: (i) haplogroup G, with 1016G and/or 989P linked to intron type-A; (ii) haplogroup L, with 1534L linked to intron-A; and (iii) haplogroup C, with 1534C and/or 1520I linked to intron-B (Figure 3).

**Table 3.**
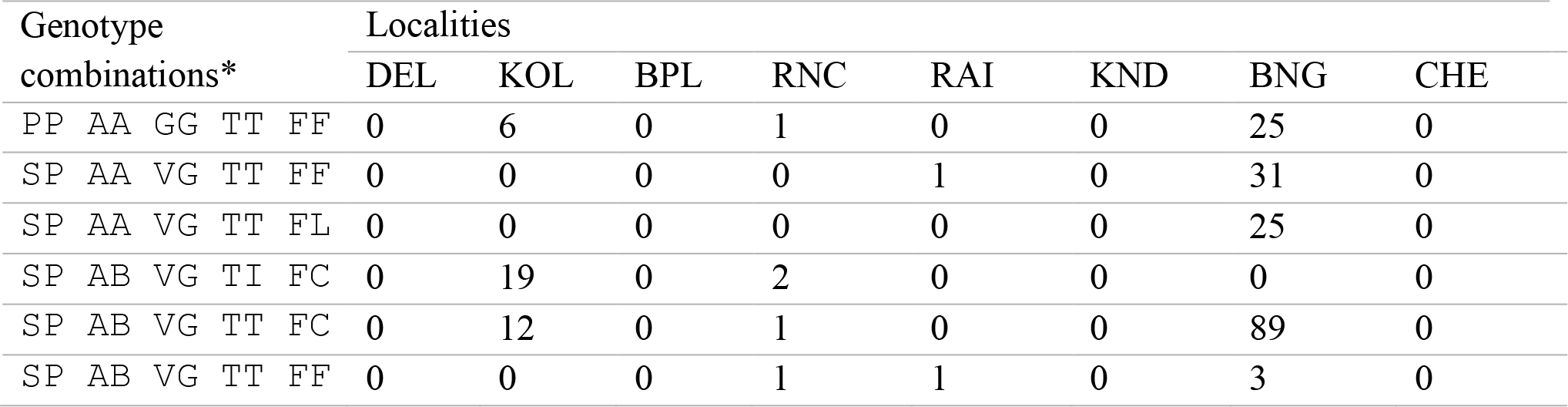
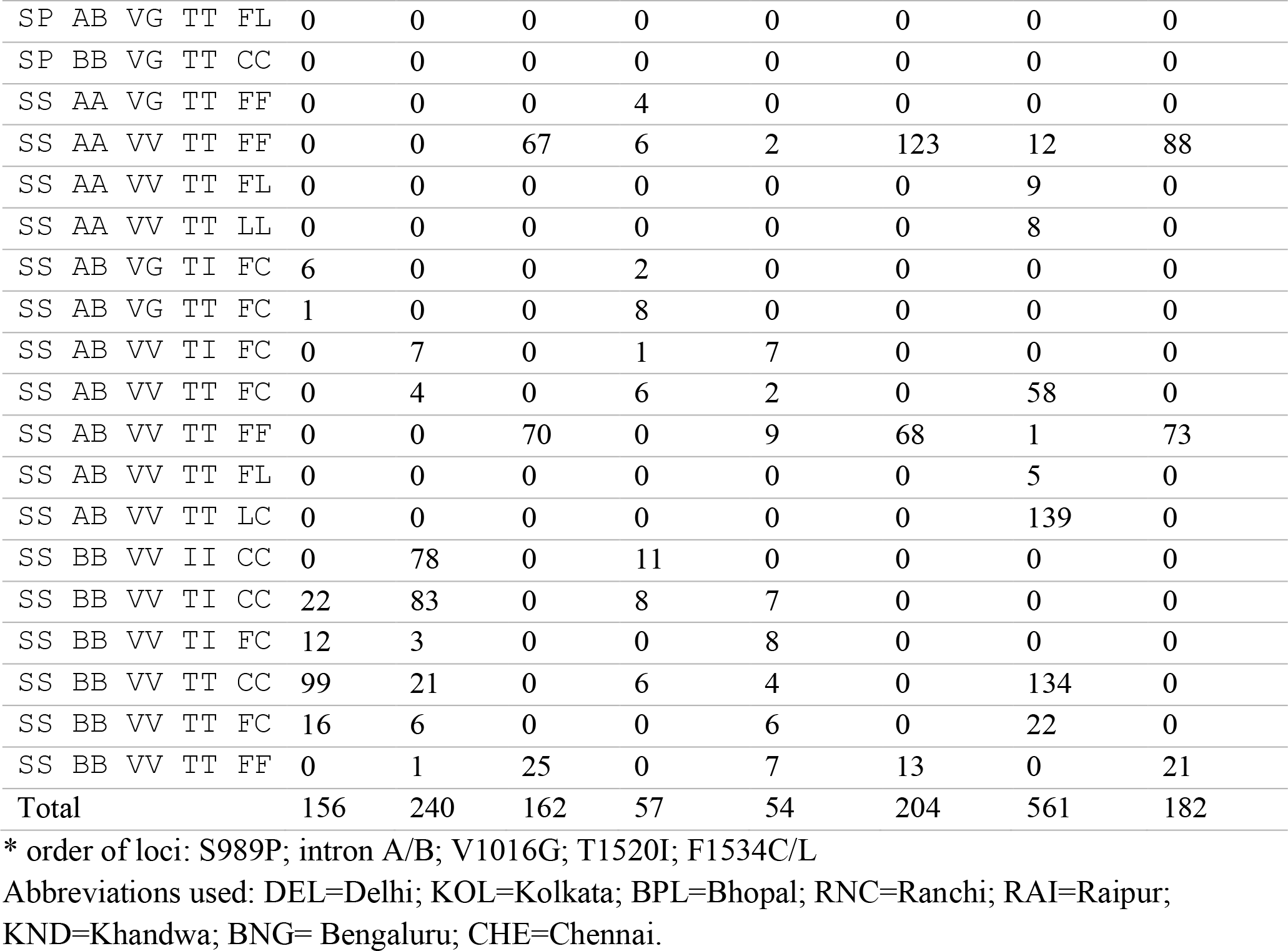
Numbers of samples with different combinations of *kdr*-genotypes and intron types in different populations.

**Table 4.** Haplotype definitions and their frequencies based on the Gibbs sampling strategy.

| S. No. | Haplotypes | Total |
| --- | --- | --- |
| 1 | SBVTC | 1018 |
| 2 | SAVTF | 947 |
| 3 | SBVTF | 438 |
| 4 | SBVIC | 365 |
| 5 | PAGTF | 249 |
| 6 | SAVTL | 194 |
| 7 | SAGTF | 21 |
|  | Total | 3232 |

**Figure 3.**
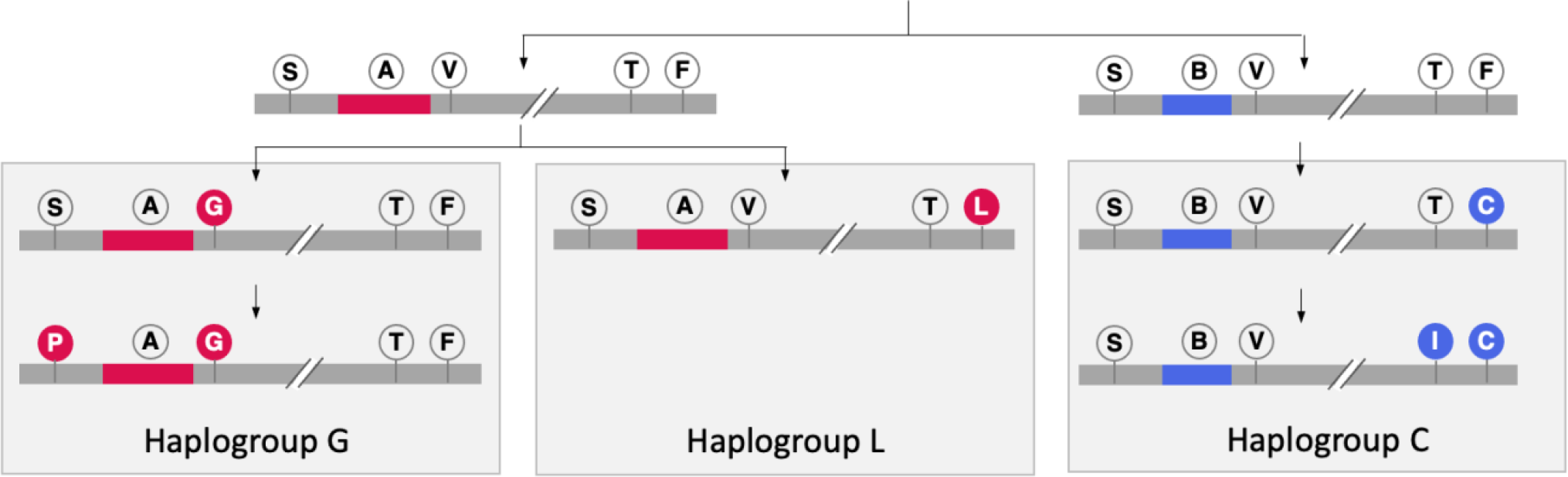
Hypothetical model illustrating the evolution of the three haplogroups (G, L, and C) based on *kdr* mutations and intron types. The 989P and 1520I mutations, occurring at lower frequencies than 1016G and 1534C, are assumed to have emerged later. (Abbreviation used: S=S989, P=989P, A=intron type A, B= intron type B, V=V1016, G=1016G, T=T1520, I=1520I, F=F1534, C=1534C, L=1534L)

Interestingly, the F1534C mutation, which is linked to intron-B, exhibits a markedly high frequency among intron-B haplotypes. In Kolkata, Ranchi, Delhi, and Bengaluru, the frequency of F1534C among these haplotypes is 97%, 99%, 91%, and 95%, respectively, suggesting that this mutation is nearing fixation in these populations. Notably, in Delhi, the V1016G mutation (without S989P) was recorded for the first time at a low frequency (2%), exclusively linked with the intron-A. Remarkably, all instances of intron-A carried the V1016G mutation, demonstrating a 100% allelic frequency within this haplotype group.

## Discussion

This study provides a comprehensive analysis of *kdr* mutations in *Ae. aegypti* populations across different regions of India, with an emphasis on understanding the prevalence and distribution of these mutations and their linkage association including intron haplotypes. The key findings indicate the presence of five *kdr* mutations (S989P, V1016G, T1520I, F1534C, and F1534L) with varying geographical distributions and frequencies. Furthermore, two distinct and conserved intron haplotypes (A and B) were identified between exons 20 and 21 of the voltage-gated sodium channel (VGSC) gene, which show strong linkage associations between specific *kdr* mutations and intron haplotypes in Indian *Ae. aegypti* populations.

Our findings highlight important geographical variations in the distribution of *kdr* mutations in Indian *Ae. aegypti* populations. The most prevalent mutation, F1534C, was detected across multiple locations, including Delhi, Ranchi, Raipur, Kolkata, and Bengaluru, with allelic frequencies ranging from 51% to 89%. The T1520I mutation, which is linked with F1534C, was present in all these localities where F1534C is prevalent, except Bengaluru, suggesting potential localized selection pressures or founder effects. The V1016G mutation, which is widely reported in Southeast Asian countries, was found in low frequencies in five Indian populations, i.e., Ranchi, Raipur, Kolkata, Bengaluru and Delhi, with the S989P mutation, typically associated with V1016G, except in Delhi.

The S989P and V1016G mutations were consistently linked with intron type A, while the F1534C mutation was linked with intron type B. This finding is consistent with earlier studies from other Asian countries, where similar associations have been documented [8, 11, 31, 36]. We identified three major haplogroups based on *kdr* mutations and intron types, i.e., haplogroup-G (V1016G with/or without S989P and with intron-A), -L (F1534L and intron-A), and -C (F1534C with/or without T1520I and with intron-B). The absence of recombination among the three haplogroups in Indian populations suggests that these mutations are maintained within distinct genetic backgrounds, which could have important implications for the spread of high-level insecticide resistance.

The co-occurrence of 989P, 1016G and 1534C mutations is very common in Asian countries [6. 37, 40] but 1016G (with or without 989P) and 1534C are present on different haplotypes. Presence of these *kdr* mutations, i.e., 989P+1016G+1534C on same haplotype has been shown to reduce sensitivity to permethrin by 1100-fold and to deltamethrin by 90-fold with resultant potential that a single recombination could result in extreme resistance [26]. This underscores the potential challenges in controlling *Aedes aegypti* populations in India using pyrethroid-based interventions. However, the absence of triple mutations in our study could suggest that while significant resistance is present, it may not yet be at the critical level that would compromise current vector control strategies. Several studies have reported co-occurrence of these triple mutations (989P+1016G+1534C) on the same chromosome but only in a few individuals in Saudi Arabia [14, 38, 39] Malaysia [40] Myanmar [37] and Indonesia [41]. In Beijing, China, all the three *Ae. aegypti* mosquitoes collected in a study, were homozygous for 989P and 1016G and heterozygous for 1534C [42]. Kasai et al. [8], also observed relatively high frequencies of V1016G + F1534C mutations (without S898P) on single haplotypes in various populations: 4.1% in Hanoi, 11.8-16.5% in Dak Lak, and 18.6% in Phnom Penh. Identifying intron types would be useful to determine whether the haplotype with the F1534C+V1016G mutations in the study areas (Vietnam and Cambodia) is due to the independent origin of either mutation or is a result of recombination. Overall, these findings indicate different evolutionary pressures or gene flow dynamics between these regions. The absence of recombination between the major haplogroups (G, L, and C) bearing *kdr* alleles in Indian populations further suggests that these mutations have been maintained separately, potentially due to strong linkage disequilibrium or limited genetic exchange. The absence of triple mutations (S989P, V1016G, F1534C on a single haplotype) in Indian populations in this report aligns with similar findings from Taiwan [23] India [7] and Mauritania [43].where these three mutations co-occurred but lack evidence of being present on the same chromosome.

Earlier records shows the absence of V1016G *kdr*-mutation in Delhi [3], however, in this study we found seven samples from Delhi having this mutation in heterozygous condition each without 989P (six with 1534C and one with F1534). Interestingly, in Delhi, only these seven samples had intron type A in heterozygous condition and all other samples had intron B with allele frequency of 98%.

The mutations S989P, V1016G/I, and I1101M are typically linked with intron type A, while F1534C is generally associated with intron type B [23, 29, 38]. Our findings align with this pattern; however, in Ghana, the F1534C *kdr* mutation in *Ae. aegypti* has been found to be associated with intron type A [23]. In the study by Fan et al. [36] it was observed that the F1534C mutation appeared in at least two distinct haplotypes of the VGSC gene, corresponding to different clades (intron types A and B) and geographic locations. This suggests that the F1534C mutation has two independent origins, likely due to separate selective pressures and subsequent spread in different populations. Therefore, it is worth investigating whether the occurrence of the triple mutation (989P + 1016G + 1534C) on the same haplotype is due to recombination between haplotypes 989P + 1016G + intron-A + F1534 and S989 + V1016 + intron-B + 1534C, or if it is a result of an independent origin of F1534C. Genotyping intron types in conjunction with *kdr* genotypes could provide valuable insights into the evolutionary history of these mutations. Identifying intron types using DNA sequencing is limited by the inability to resolve heterozygous sequences due to the presence of indels, which causes sequence collapse. The PCR method developed in this study for genotyping intron haplotypes can help address this issue. Also, it is crucial to sequence the wider genomic region encompassing the loci S989P, V1016G, and F1534C/L, either by amplifying through long-read PCR [44] or using single molecule next-generation sequencing. Via this approach genetic signatures of recombination events, particularly within intronic regions where recombination hotspots are predominantly located [45-46] can be identified.

In this study, sequencing a limited number of samples identified two distinct intron types, A and B. Despite their divergence, each intron type remained highly conserved, with no observed variability, regardless of the presence or absence of *kdr* mutations. Intron-A matched the sequences found in haplotypes V4, V8, and V9 (have identical intron), while intron-B matched to V2 haplotype, as defined by Fan et al. [36]. Fan et al. reported that the 989P and 1016G mutations are linked to intron-A, which is consistent with our findings. They also observed an association between the 1534C mutation and V2 (intron-B) in Asian populations, similar to our results, with the exception of one sample (out of 174 samples, excluding those from the Americas and Africa).

The F1534C mutation was found exclusively in locations where intron type B was the predominant haplotype. Furthermore, in several areas, including Kolkata, Ranchi, Delhi, and Bengaluru, the F1534C mutation is nearly fixed among intron-B haplotypes, with frequencies ranging from 91% to 99%. In contrast, a similarly high frequency of the 1016G mutation among intron-A haplotypes was observed only in Delhi, where all seven samples with intron-A (in heterozygous condition) carried this mutation.

This study has some limitations. The geographic coverage was restricted to specific regions, and further research is needed to evaluate the prevalence of *kdr* mutations in other parts of India. Additionally, functional studies on mutations like S989P and T1520I that are linked to V1016G and F1534C, respectively, are needed to explore the potential compensatory effects of these mutations on mosquito fitness or their additive role in conferring resistance phenotype, which could provide further insights into the evolutionary dynamics of resistance. Finally, there is a need to investigate other resistance mechanisms.

In conclusion, our study provides a detailed characterization of *kdr* mutations and their genetic associations in Indian *Ae. aegypti* populations. While integrated vector management approaches, combining chemical, biological, and environmental control methods, are necessary, an immediate priority should be to use of pyrethroids cautiously in populations where *kdr* mutations are prevalent. In areas without documented *kdr* mutations, resistance profiling should be conducted to assess whether pyrethroids might still be a viable option in the short term. Continued surveillance and targeted interventions will be essential for maintaining the effectiveness of vector control programs and reducing the burden of arboviral diseases in India.

## Acknowledgements

We are grateful to staff namely Mr. Uday Prakash, Kanwar Singh, Bhupal Arya for their excellent technical assistance rendered for DNA isolation and PCR amplifications.

